# Normal Uterine Fibroblast Are Reprogramed into Ovarian Cancer-Associated Fibroblasts by Ovarian Tumor-derived Conditioned Media

**DOI:** 10.1101/2023.09.29.560158

**Authors:** Hailey Axemaker, Simona Plesselova, Kristin Calar, Megan Jorgensen, Jared Wollman, Pilar de la Puente

**Affiliations:** Cancer Biology and Immunotherapies Group, Sanford Research, Sioux Falls, SD 57104, USA; MD/PhD Program, University of South Dakota Sanford School of Medicine, Sioux Falls, SD 57105, USA; Flow Cytometry Core, Sanford Research, Sioux Falls, SD 57104, USA; Department of Obstetrics and Gynecology, University of South Dakota Sanford School of Medicine, Sioux Falls, SD 57105, USA; Department of Surgery, University of South Dakota Sanford School of Medicine, Sioux Falls, SD 57105, USA

## Abstract

Cancer-associated fibroblasts (CAFs) are key contributors to ovarian cancer (OC) progression and therapeutic resistance through dysregulation of the extracellular matrix (ECM). CAFs are a heterogenous population derived from different cell types through activation and reprogramming. Current studies rely on uncharacterized heterogenous primary CAFs or normal fibroblasts that fail to recapitulate CAF-like tumor behavior. Here, we present a translatable-based approach for the reprogramming of normal uterine fibroblasts into ovarian CAFs using ovarian tumor-derived conditioned media to establish two well-characterized ovarian conditioned CAF lines. Phenotypic and functional characterization demonstrated that the conditioned CAFs expressed a CAF-like phenotype, strengthened proliferation, secretory, contractility, and ECM remodeling properties when compared to resting normal fibroblasts, consistent with an activated fibroblast status. Moreover, conditioned CAFs significantly enhanced drug resistance and tumor progression and resembled a CAF-like subtype associated with worse prognosis. The present study provides a reproducible, cost-effective, and clinically relevant protocol to reprogram normal fibroblasts into CAFs using tumor-derived conditioned media. Using these resources, further development of therapeutics that possess potentiality and specificity towards CAF-mediated chemoresistance in OC are further warranted.

## INTRODUCTION

Ovarian cancer (OC) is one of the deadliest forms of cancer in women, where more than 80% of patients develop chemotherapy resistance, resulting in advanced recurrence and eventually death ^1^. It is usually diagnosed at a late stage and a high tumor grade, with a very low rate of women being diagnosed at an early stage. Treatment barriers in ovarian cancer have been increasingly attributed to stromal reprogramming and activation, extracellular matrix (ECM) remodeling, angiogenesis, and drug delivery ^1–3^. A known source for these barriers is stromal cell types, which are a main contributor to hallmarks of cancer like tumor recurrence, metastasis, and chemotherapy resistance ^3–5^. There is evidence that stromal cells are recruited by the tumor microenvironment (TME) through tumor and stromal cell crosstalk, supplying tumor-associated stromal cells to a developing tumor, and contributing to its proliferation ^4,6^. Stromal cells are the main secretors of the ECM providing a structural framework of connective tissue throughout the body. In tumors, the most important type of stromal cells are cancer-associated fibroblasts (CAFs), a type of activated fibroblast.

CAFs may be derived from several cell types including fibroblasts, mesenchymal stem cells (MSCs), or epithelial and endothelial cells through activation and reprogramming by cancer cells, epithelial to mesenchymal transition (EMT), and endothelial to mesenchymal transition (EndMT), respectively ^7^. Stromal cell types are recruited by chemokines, cytokines, and growth factors to specific cancer sites and the dysregulated ECM at these sites activates them into CAFs. These different sources of activation lead to heterogeneity of CAFs and a variety of different subtypes, many of which have not been clearly identified yet. CAFs are key contributors to tumor progression and therapeutic resistance through the remodeling of the tumor ECM composition and structure ^8^. An example of this is CAFs ability to induce EMT through the upregulation of transforming growth factor beta (TGF-β) ^9^. CAFs are inflammatory and are mainly known to be pro-tumorigenic, largely attributed to their production of cytokines like interleukin-6 (IL-6) and CC-chemokine ligand 2 (CCL2/MCP-1) that interfere with T cell function ^10–12^. Thus, CAFs are considered to be notable players in promoting many aspects of tumor function and a promising targeted therapeutic option.

Current models studying the stromal cell influence on OC rely heavily on normal fibroblasts including NIH3T3 fibroblasts, non-ovarian tissue origin such as dermal and lung fibroblasts, or immortalized ovarian fibroblasts (TRS3 and NOF151-hTERT) ^13–16^. Unfortunately, the effects of CAFs and normal fibroblasts in OC progression and drug resistance are significantly different ^17–20^, making normal fibroblasts a less ideal source. Alternatively, primary CAFs can be isolated from tumors and cultured *in vitro*, but their lifespan is finite; they have shown limited expansion capacity and the use of cells from different patients can result in non-reproducible results associated with the technologies and methodologies from each laboratory, and the lack of characterization, and heterogeneity between patients ^21–23^. Therefore, there is a need to establish protocols that allow for the reprogramming of normal fibroblasts into CAFs with a reproducible, cost-effective, and clinically relevant approach.

The overall objective of this investigation is to present a translatable-based approach to reprogram human uterine fibroblasts (HUFs) into CAFs using ovarian tumor-derived conditioned media, followed by phenotypic and functional characterization of the conditioned CAFs. Our results demonstrate the importance of activated CAFs on several OC processes, providing a protocol and two well-characterized ovarian conditioned CAFs to study the effect of the OC tumor microenvironment (TME). In the absence of reproducible and functionally characterized protocols to engineer and reprogram CAFs in culture, the development of effective anticancer therapies that mitigate CAF-mediated chemotherapy resistance in OC will likely remain difficult.

## RESULTS

### Conditioned CAFs Express CAF-Like and Stromal Markers

To phenotypically characterize our conditoned CAFs, we evaluated their cell morphology and surface marker expression. All stromal cells including HUFs, conditioned CAFs, and primary OC-CAFs were monitored routinely for morphological changes over passages. The cells retained their fibroblast-like morphology in culture over passages (Figure 1A) and no morphological changes were found. Cell surface marker expression was assessed for all the stromal cell types, as well as the OC cell lines at early passages (P2-P4) given the limited expansion capacity of the primary CAFs. HUFs, primary OC-CAFs, both conditioned CAFs, and the adipose MSCs expressed CAF markers FAP and CD29, which are indicators of activated fibroblasts ^24,25^. HUFs, KURA-CAFs, OC-CAFs, and adipose MSCs showed negative PDGFRα expression, while SKOV-CAFs exhibited low expression. HUFs and adipose MSCs were negative for α-SMA, while both CM-CAFs and OC-CAFs display low expression (Figure 1B). HUFs, primary OC-CAFs, both conditioned CAFs, and the adipose MSCs expressed stromal markers CD90 and CD73, which are classical MSC markers ^26^, and vimentin, which is highly expressed by various types of fibroblasts ^27^. All stromal cell types remained negative for the epithelial marker EpCAM/CD326 and the immune marker CD45 (Figure S1A). KURAMOCHI and SKOV-3 OC cell lines were validated for negative FAP expression and positive CD326 expression (Figure S1B). FAP expression was also evaluated in the conditioned CAFs over passages and confirmed that FAP expression was retained in the later passages up to at least P9 (Figure S1C).

**Figure 1.**
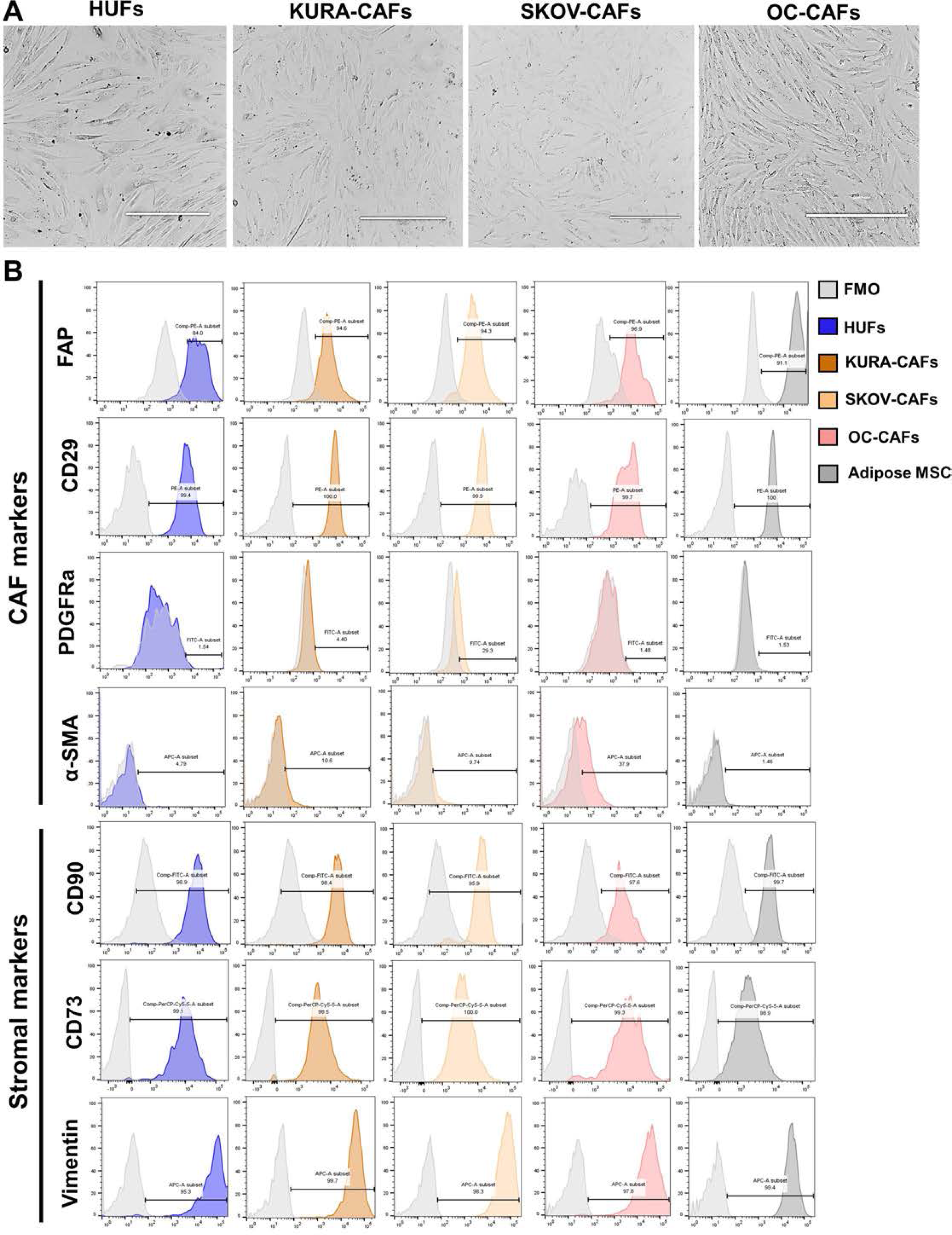
Conditioned cancer-associated fibroblasts (CM-CAFs) express CAF-like and stromal markers. (A) Representative images of 2D cultures of HUFs, KURA-CAFs, SKOV-CAFs, and OC-CAFs, scale bar = 400µm (B) Representative flow cytometry histograms of expression of CAF-related markers (FAP, CD29, PDGFRα, α-SMA), stromal markers (CD90, CD73, Vimentin), and respective fluorescence minus one (FMO) controls in HUFs, KURA-CAFs, SKOV-CAFs, OC-CAFs, and adipose-derived MSCs. Cells from early passages 2-4.

### Conditioned CAFs Possess Enhanced Expansion Capacity, a CAF-Like Cytokine Profile, and Deposit ECM as Activated Fibroblasts

To demonstrate the activation of our conditioned CAFs, we performed experiments that measured cell proliferation, cytokine secretion, and ECM deposition in HUFs, both conditioned CAFs, and primary OC-CAFs. While HUFs and OC-CAFs showed limited expansion of less than 2-fold and for limited passages (less than P4), conditioned CAFs showed significantly enhanced expansion capacity (2-4 fold) compared to primary HUFs and OC-CAFs, as well as expanded proliferation over longer passages (P7) (Figure 2A). In order to account for the potential influence of ovarian media as the source of activation, we exposed HUFs to ovarian media and it did not enhance or prolong their expansion capacity compared to HUFs in their original media (Figure 2A). Cytokine secretion profile was assesed in all stromal lines, as well as their culture media. IL-6, IL-8, MCP-1, osteprotegerin (OPG), TIMP-2 and PLGF were the most highly secreted chemokines/cytokines by the 4 cell types (Figure 2Bi and Figure S2A), although part of IL-6, IL-8, MCP-1, and TIMP-2 in the conditioned CAFs was derived from the cancer conditioned media (Figure S2A and S2B). Importantly, CM-CAFs secreted similar levels to HUFs and primary OC-CAFs of revelant tumor-promoting cytokines/chemokines including TIMP-1, CXCL5, GRO, VEGF, and TGF-β (Figure 2Bii). The cytokines/chemokines IL-6, IL-8, and MCP-1 are known to function as tumor promotors while also supporting the migration of OC cells ^28–30^. Specifically, TGF-β, is a master regulator of fibrosis and a major secreted factor by CAFs; GRO, is an IL-8 related cytokine ^31^, and TIMP-1 and TIMP-2, are both important tissue inhibitors of metalloproteinases governing ECM remodeling ^32^. OC-CAFs were the only cell type showing high expression of LIF, but it should be noted that part of it is coming from their different culture media (Figure 2Bii and Figures S2A and S2B). Collagen type I deposition was assesed by immunohistochemistry to demonstrate that conditioned CAFs deposited ECM like activated fibroblasts. Figure 2C shows that there is higher collagen positivity in both conditioned CAFs than in HUFs, confirming their fibroblast activation.

**Figure 2.**
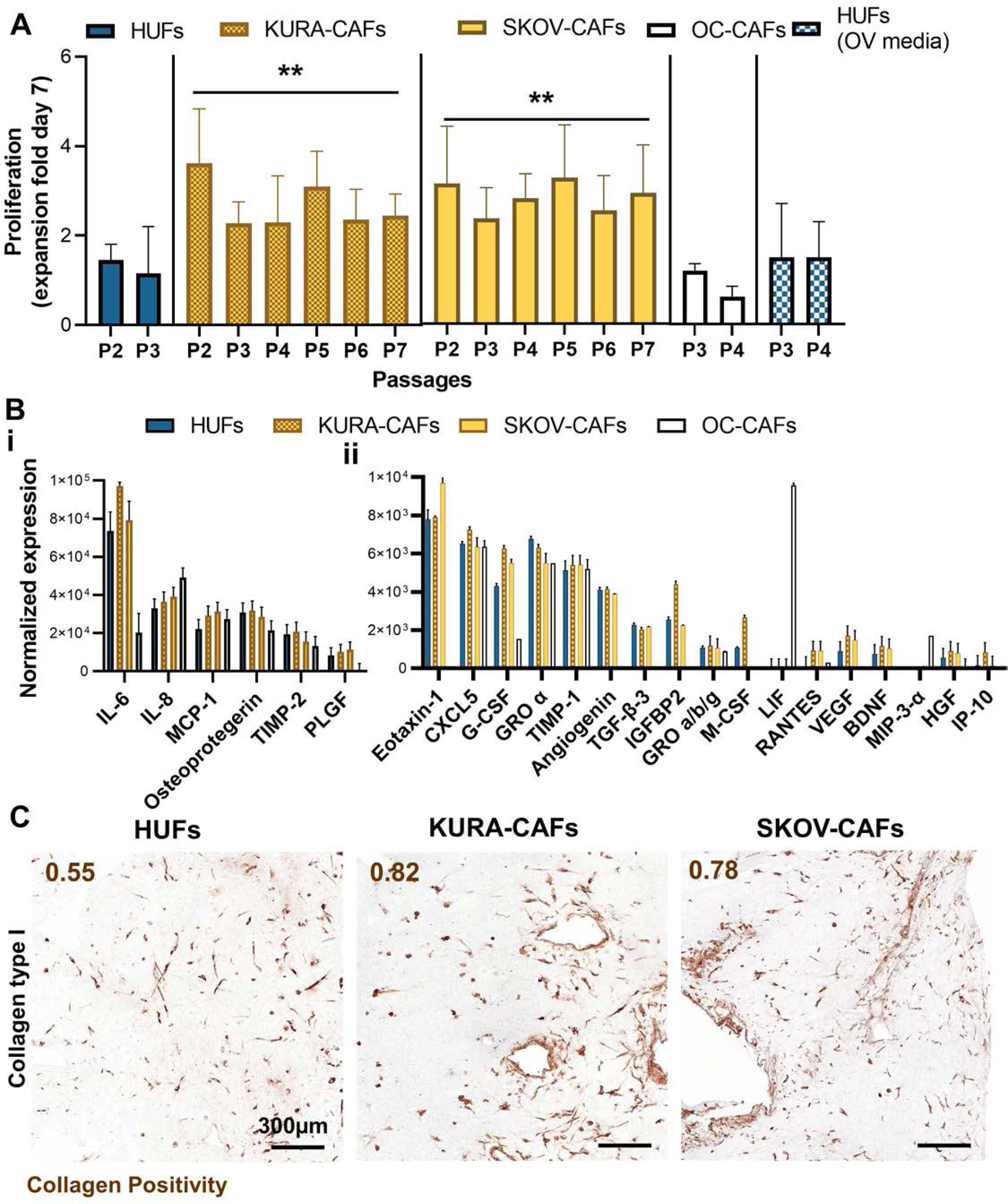
Conditioned CAFs possess enhanced expansion capacity, CAF-like cytokine profile, and deposit ECM as activated fibroblasts. (A) Proliferation of HUFs, KURA-CAFs, SKOV-CAFs, and OC-CAFs in culture over passages presented as the expansion fold after 7 days for each passage; HUFs were also cultured in ovarian (OV) media (mean ± SD, n = 4). (B) Normalized expression of cytokines expressed in media cultured for 4 days with HUFs, KURA-CAFs, SKOV-CAFs, and OC-CAFs, respectively, where i) represents high cytokine expression and ii) moderate cytokine expression. (C) Representative IHC images of HUFs, KURA-CAFs, and SKOV-CAFs for Collagen type I expression on day 7 with quantified positivity (Npositive/Ntotal), scale bar = 300μm. **p>0.001

### Conditioned CAFs Are Implicated in Contractility, Tumor Promotion, and Drug Resistance

Collagen gel contractility assays, tumor progression and drug resistance in 3D co-cultures, ALDH expression, and spheroid formation were assessed to functionally characterize the conditioned CAFs. HUFs, both conditioned CAFs, and primary OC-CAFs were examined for collagen contractility capabilities by measuring the percent change of the area of collagen 3D matrices over 24 hours as an indication of fibroblast activation. While HUFs produced a very slight contraction of collagen gels of about 20%, primary OC-CAFs and conditioned CAFs contracted significantly more compared to HUFs with about 50% and 60-70% contraction, respectively (Figure 3A). CAFs have shown clear roles in tumor promotion and induction of drug resistance, in order to assess these mechanisms, physiologically relevant 3D models allowing cell-cell and cell-ECM interactions were used. Co-culture of cancer and conditioned CAFs or OC-CAFs enhanced cancer growth significantly about 2.5-3-fold compared to both cancer alone and cancer in 3D co-culture with HUFs for both cancer lines (KURAMOCHI and SKOV-3) (Figure 3B). Similarly, 3D co-cultures of cancer and corresponding conditioned-CAFs significantly reduced cell killing by carboplatin treatment when compared to cancer alone or cancer co-cultured with HUFs for both lines, and have no significant differences with primary OC-CAFs (Figure 3C). ALDH activity was assessed by aldefluor expression and conditioned CAFs and OC-CAFs had significantly higher ALDH+ cells than HUFs (Figure 3D and *3E*). Finally, spheroid formation, integrity and composition were assessed in co-culture conditions. Conditioned CAFs and OC-CAFs produced spheroids with significantly greater area, length, and integrity than when cancer cells were co-cultured with HUFs (Figure 3F, 3G and Figure *S*3A). After digestion, spheroids showed similar epithelial/stromal distribution when compared to the co-cultures with the different stromal types, although there is a slight trend of higher epithelial composition with conditioned CAFs and OC-CAFs compared to co-cultures with HUFs for both OC lines (Figure *S*3B and *3B*C). Finally, we assessed whether the reprograming of HUFs with conditioned cancer media could affect the conditioned CAFs tumorigenic potential. Neither HUFs, conditioned CAFs, nor OC-CAFs induce tumors *in vivo* when implanted in immunodeficient mice. Mice had a normal weight gain over 6 weeks and no tumors were palpable (Figure S3D).

**Figure 3.**
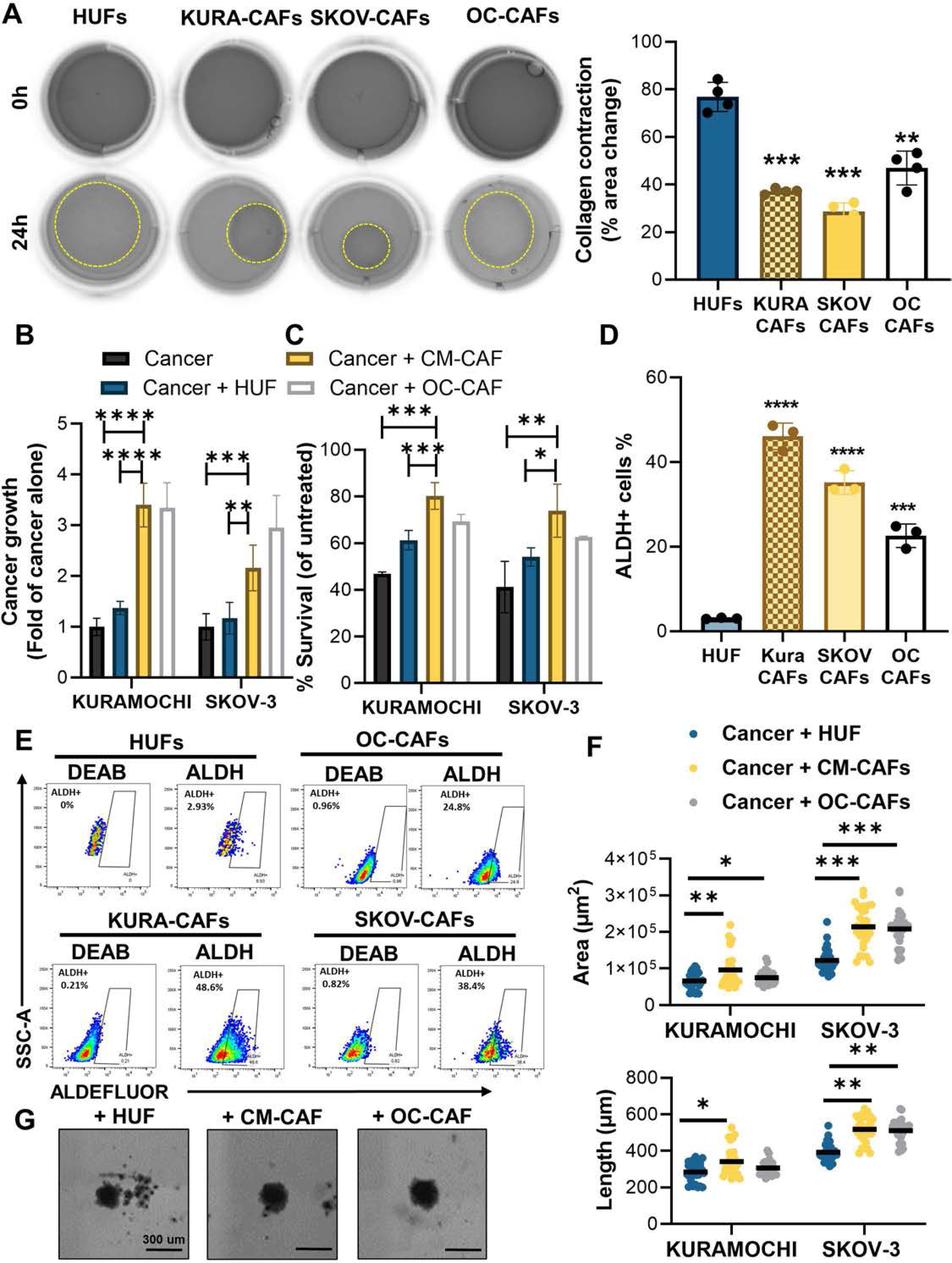
Conditioned CAFs are implicated in contractility, tumor promotion and drug resistance. (A) Representative images of collagen contractility assay for HUFs, KURA-CAFs, SKOV-CAFs, and OC-CAFs at 0 hours and 24 hours. Collagen contraction is denoted as percentage of area change after 24 hours. (B) Effect of stromal cells (HUF, CM-CAF, OC-CAF) on cancer growth (KURAMOCHI, SKOV3) where cancer was cultured in a 3D matrix for 7 days either alone or in co-culture with HUF, CM-CAF, or OC-CAF, respectively; shown as the fold of cancer alone (n = 3). (C) Effect of carboplatin (30uM) on cancer survival (KURAMOCHI, SKOV3) cultured in 3D matrix either alone or in co-culture with stromal cells (HUF, CM-CAF, OC-CAF) for 7 days, normalized to untreated (PBS control) (n = 3). (D) ALDH activity shown as percentage of ALDH positive cells in HUFs, KURA-CAFs, SKOV-CAFs, and OC-CAFs. (E) Representative flow cytometry images showing gating of ALDH positive cells by comparing aldefluor stained cells to cells stained with the DEAB control. (F) Effect of stromal cells (HUF, CM-CAF, OC-CAF) on cancer (KURAMOCHI, SKOV3) spheroid formation in culture where cancer was co-cultured with respective stromal cells and grown for 21 days by comparing spheroid length and area measurements. (G) Representative Cytation3 images of day 21 KURAMOCHI cancer spheroids co-cultured with HUFs, KURA-CAFs, and OC-CAFs, respectively. Scale bar = 300μm. Results are shown as mean ± standard deviation and were analyzed using ANOVA where *p>0.05, **p>0.001, ***p>0.0001, ****p>0.00001.

### Conditioned CAFs Reveal a CAF-like transcriptional signature with involvement of the ECM/matrisome

To understand how tumor conditioned media alters the transcriptome of the CM-CAFs, we compared the transcriptional profiles of HUFs, CM-CAFs and primary OC-CAFs by bulk RNA sequencing. We found 748, 790, and 1543 differentially expressed genes (DEG) when KURA-CAFS, SKOV-CAFs, and primary OC-CAFs were compared to HUFs respectively (Figure 4A). To identify pathways enriched by the DEGs in conditioned CAFs and OC-CAFs, gene ontology analysis revealed ECM-related pathways (extracellular matrix structural constituent, extracellular matrix binding, laminin binding, integrin binding and growth factor activity) to be similarly enriched in both conditioned CAFs and OC-CAFs when compared to HUFs, pathways relevant to provide structural support to the ECM ^33,34^ (Figure 4B). When the DEGs were compared we found a 29%, 28%, and 14% overlap of the genes for KURA-CAFs, SKOV-CAFs, and OC-CAFs respectively (Figure 4Ci). Significantly, out of the total DEGs produced, 84, 92, and 186 belonged to the matrisome, an ensemble of genes encoding for ECM-related genes, clearly highlighting the implication of CM-CAFs and OC-CAFs in the matrisome/ECM. Importantly, mesenchymal subtypes characterized by CAF presence have been linked to a worse outcome in OC ^35^. When tumors of mesenchymal subtype were compared to HUFs and the generated DEGs were compared to the DEGs from CM-CAFs to HUFs, we found that a total of 371 overlapped genes accounted for 60% of the total DEGs for the CM-CAFs, clearly highlighting the relevance of the CM-CAFs-like transcriptional profile in patient outcomes (Figure 4Cii).

**Figure 4.**
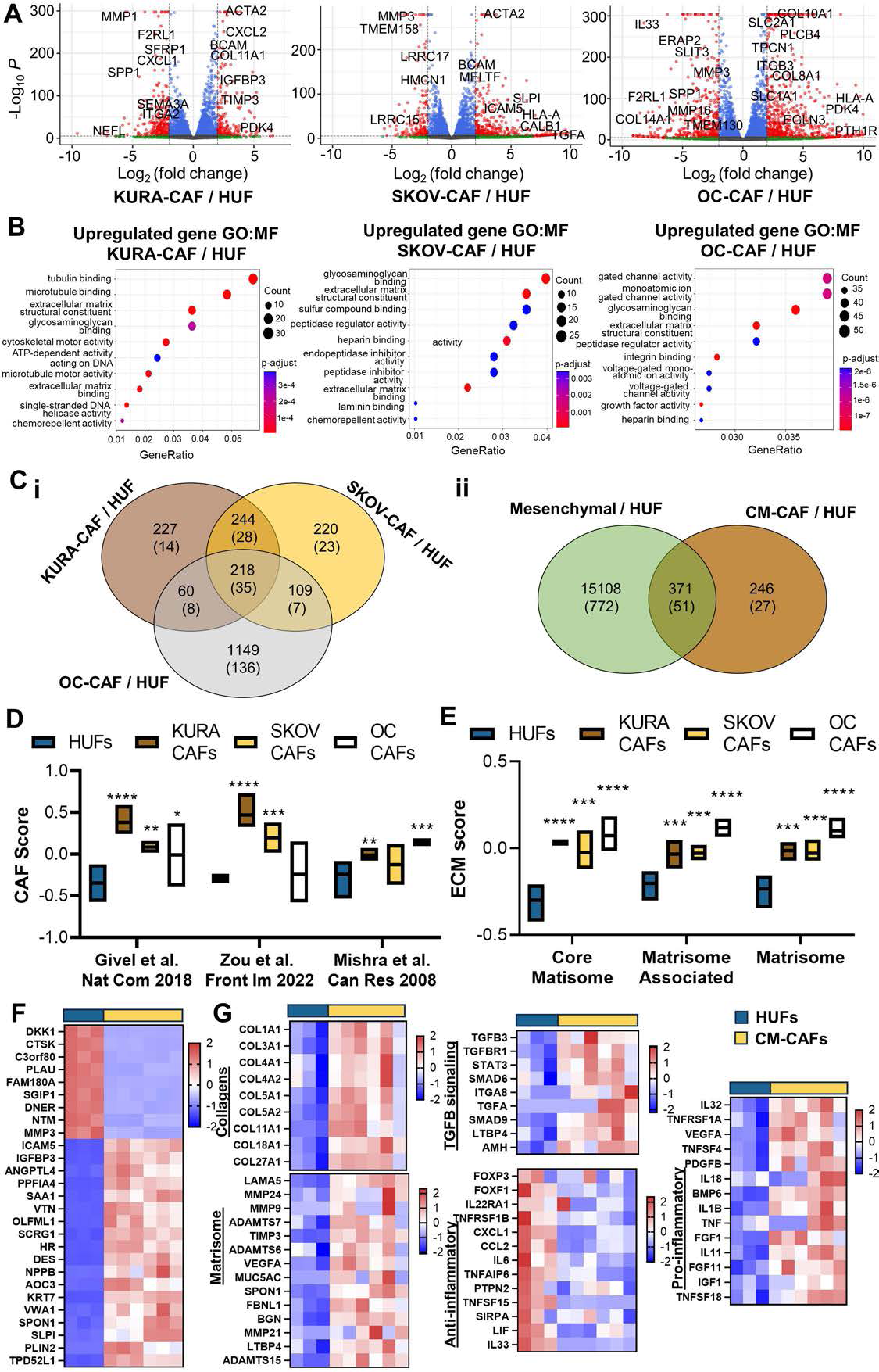
Conditioned CAFs Reveal a CAF-like transcriptional signature with involvement of the ECM/matrisome. (A) Volcano plots of significant upregulated and downregulated genes from bulk RNA-seq results comparing OC-CAFs, SKOV-CAFs, and KURA-CAFs to HUFs, respectively. Red = p-value and Log_2_ fold change, Blue = p-value, Green = Log2 fold change, and Gray = Not significant. (B) Dot plots of significantly enriched GO:MF terms in differentially expressed genes from bulk RNA-seq comparing OC-CAFs, SKOV-CAFs, and KURA-CAFs to HUFs, respectively. (C) Venn diagrams comparing the overlap of differentially expressed genes (DEGs) between i) OC-CAFs versus HUFs, SKOV-CAFs versus HUFs, and KURA-CAFs versus HUFs and between ii) DEGs from patients with mesenchymal subtype versus HUFs and CM-CAFs versus HUFs; number in parentheses denotes number of matrisome ECM-related DEGs. (D) CAF score of HUFs, KURA-CAFs, SKOV-CAFs, and OC-CAFs when scored by GSVA using three different CAF-related gene sets. E) ECM score of HUFs, KURA-CAFs, SKOV-CAFs, and OC-CAFs when scored by GSVA using three different sets of matrisome ECM-related genes, respectively. (F) Heatmap of differentially expressed genes regulated by reprogramming of normal fibroblasts into CAFs. (G) Heatmaps of differentially expressed genes of KURA-CAF and SKOV-CAF (CM-CAFs) versus HUFs related to collagens, matrisome, TGF-beta (TGFB) signaling, anti-inflammatory activity, and pro-inflammatory activity. Results are shown as the mean and were analyzed using ANOVA where *p>0.05, **p>0.001, ***p>0.0001, ****p>0.00001.

Moreover, the stromal cells were ranked according to three distinct CAF scores ^36–38^ and robustly showed that both CM-CAFs and OC-CAFs significantly ranked higher for CAF scores than HUFs (Figure 4D). Similarly, considering the clear implication of CAFs in ECM remodeling, three ECM scores were produced ^39,40^, and again both CM-CAFs and OC-CAFs significantly ranked higher for ECM scores than HUFs (Figure 4E). Comparably, some of the DEGs that significantly regulate HUFs reprograming into CAFs are matrisome/ECM related including IGFBP3, ANGLPTL4, SPON1, SLPI, VTN and VWA1 (Figure 4F) ^41–45^. Conditioned media induced profound changes in the transcriptome of the HUFs, particularly in genes significantly enriched as part of pathways, such as growth factors like TGF-β, collagens, the ECM/matrisome, and immunomodulatory cytokines, and as anticipated we see upregulation of these genes in the CM-CAFs and downregulation in the HUFs (Figure 4G).

### Conditioned CAFs Preserve CAF Phenotype After Removal of Tumor-Derived Conditioned Media

To demonstrate that our conditioned CAFs have been transformed/reprogrammed into CAFs and that they remain CAF-like when removed from tumor-derived conditioned media, we conducted retrieval experiments, comparing CAFs kept in the conditioned media to CAFs switched to ovarian media. Conditioned CAFs at post-retrieval passages P1, P3 and P5 (R-P1, R-P3, and R-P5) showed the same contractility potential as conditioned CAFs (Figure 5A). Moreover, the retrieval CAFs promoted growth the same as the corresponding conditioned CAFs with no changes to tumor promotion in both CM-CAFs lines (Figure 5B). Finally, we also confirmed that the classical CAF marker (FAP) was retained (Figure 5C), as well as stromal CD73 expression (Figure 5D), while epithelial CD326 expression remained negative (Figure 5E) over the 5 post-retrieval passages for both lines.

**Figure 5.**
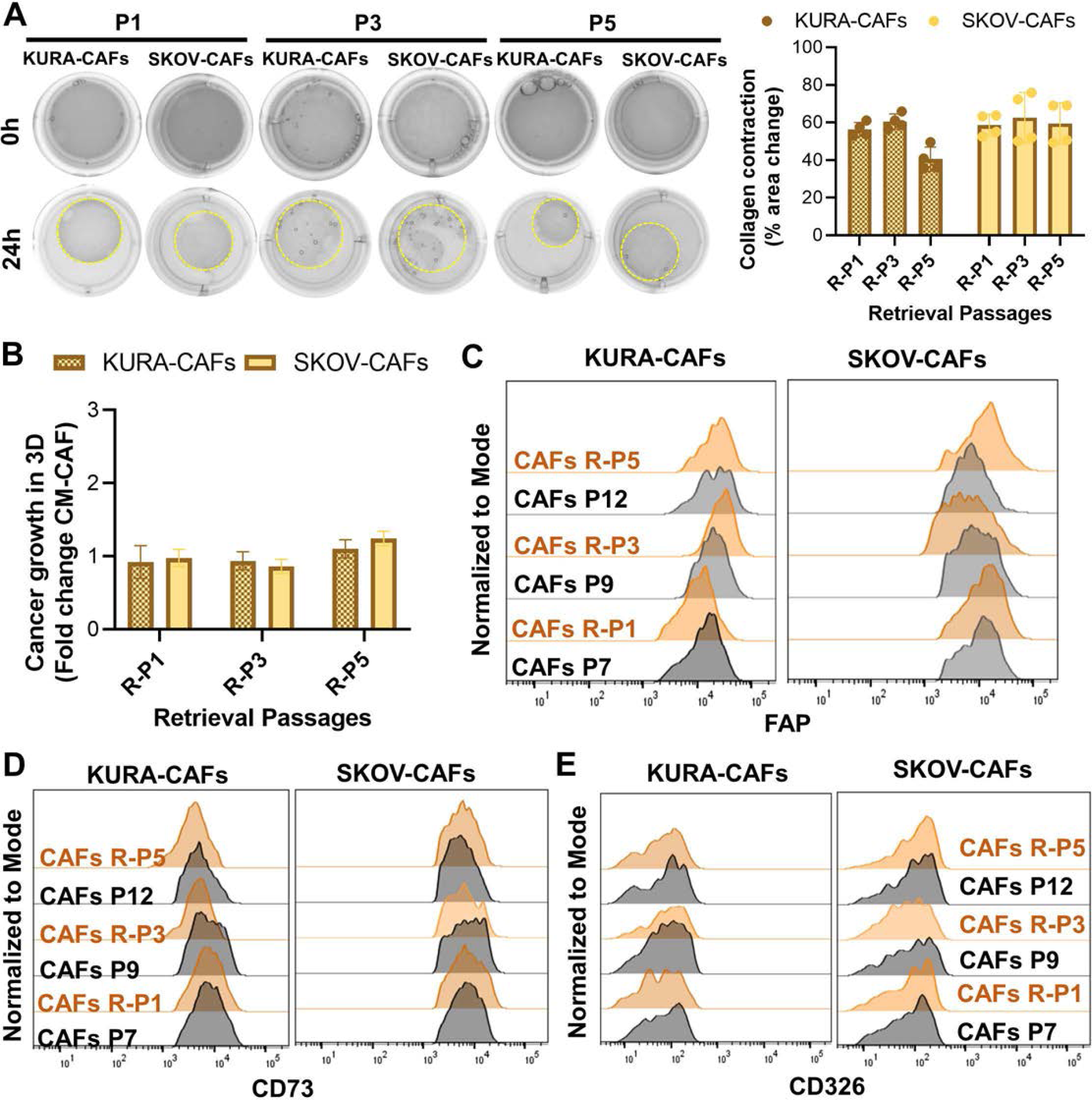
Conditioned CAFs preserve CAF phenotype after removal of tumor-derived conditioned media. (A) Representative images of collagen contractility assays at 0 hours and 24 hours for KURA-CAFs and SKOV-CAFs at post-retrieval passages P1, P3, and P5. Collagen contraction is denoted as percentage of area change after 24 hours. (B) Effect of KURA-CAFs and SKOV-CAFs at post-retrieval passages P1, P3, and P5 on cancer growth (KURAMOCHI, SKOV3) where cancer was cultured in a 3D matrix in co-culture with either the conditioned KURA-CAFs and SKOV-CAFs or the retrieval passages of KURA-CAF and SKOV-CAF, respectively, for 7 days; shown as the fold of corresponding conditioned-media CAFs (CM-CAF) (n = 3). (C) Representative flow cytometry histograms of FAP expression in conditioned KURA-CAFs and SKOV-CAFs at post-retrieval passages P1, P3, and P5 and corresponding non-retrieved CM-CAFs. (D) Representative flow cytometry histograms of stromal marker (CD73) expression in conditioned KURA-CAFs and SKOV-CAFs at post-retrieval passages P1, P3, and P5 and corresponding non-retrieved CM-CAFs. (E) Representative flow cytometry histograms of epithelial marker (CD326) expression in conditioned KURA-CAFs and SKOV-CAFs at post-retrieval passages P1, P3, and P5 and corresponding non-retrieved CM-CAFs.

## DISCUSSION

In this study, we were able to demonstrate that normal uterine fibroblasts (HUFs) can be reprogrammed into ovarian cancer associated fibroblasts (CAFs) using ovarian tumor-derived conditioned media. We have optimized a translatable-based approach to transform fibroblasts into CAFs and validated phenotypically and functionally two ovarian conditioned CAFs lines. This protocol relies only on paracrine secretion transformation, avoiding complicated co-culture techniques where cells are in physical interaction. This study demonstrates that ovarian tumor-derived conditioned media is sufficient to induce fibroblast reprogramming into CAFs, which is in line with other reports using secreted factors from conditioned media to reprogram normal cells into cancer-associated ones ^46–50^.

The approach presented in this study provides an alternative to investigations into ovarian cancer progression and chemoresistance with well-characterized ovarian CAFs, avoiding the use of normal fibroblasts, immortalized lines, or heterogenous primary sources. Importantly, the ovarian conditioned CAFs possess a CAF-like phenotype, strengthened proliferative, secretory, contractility, and ECM remodeling properties when compared to resting normal fibroblasts, as it has been previously documented ^17,51,52^, and their activated fibroblast status was demonstrated in Figures 1 to 4. Importantly, CAFs have pro-tumorigenic functions by secreting tumor-promoting signals including TGF-β and IL-6 ^53,54^, and they contribute to the development of resistant cancer phenotypes following chemotherapy by paracrine signaling via cytokines and exosomes ^51,55–57^. Unfortunately, immortalized fibroblast lines undergo genetic alterations that can create unique gene patterns that might express attributes or functions distinct to the original cells ^58^. While primary CAFs derived from tumors would be ideal in terms of the origin and activated fibroblast status, lack of characterization considering that ovarian cancer patients present wide heterogeneity, makes it challenging to broadly adopt their use. The development of characterized CAFs lines could limit some of those variables.

One predicament in defining CAFs is the lack of a pan-specific marker, or a consensus of markers to identify CAFs, adding to the difficulty in differentiating CAFs from other cell types. Consequently, phenotypic and functional characterizations are needed in order to fully characterize CAFs. To date, CAFs are defined as cells that express different levels of mesenchymal biomarkers such as FAP, α-SMA, platelet-derived growth factor receptor alpha (PDGFRα) and vimentin, and lack the expression of markers for epithelial or hematopoietic cells ^27,59^. Importantly, functional CAF subsets sustain a unique cytokine expression profile shaping the tumor microenvironment. Significant advances have been made in pancreatic cancer CAF subtyping and characterization of their different functionality ^60–64^, yet investigations in OC are limited. Recent OC studies have yielded four CAF subtypes, described as CAF S1–S4 based on the differential expression of five fibroblast markers (FAP, CD29, α-SMA, FSP1, and PDGFRβ) ^38^. Critically, a CAF-S1 subtype expressing FAP^High^ SMA^Med-High^ FSP1^Med-High^ CD29^Med-High^ PDGFRβ^Med^ was associated with worse survival. Similarly in a recent study, OC CAFs identified as FAP^High^ SMA^Low^ subpopulation were related with worse overall survival ^65^. The characterized ovarian conditioned CAFs express FAP^High^ SMA^Low^ CD29^High^ PDGFRα^Neg^ and FAP^High^ SMA^Low^ CD29^High^ PDGFRα^Low^ for KURA-CAFs and SKOV-CAFs, respectively (Figure 1). Likewise, Tothill et al. ^35^ reported four distinct subtypes in high-grade serous ovarian carcinoma, with a poor prognosis subtype displaying strong stromal response, correlating with extensive desmoplasia. When the transcriptomic profile of this mesenchymal subtype was compared to HUFs and those transcriptomic changes of the conditioned CAFs against HUFs, we found a significant 60% overlap of DEGs (Figure 4). Another study identified two subpopulations of CAFs in epithelial OC by their expression of seven CAF-related markers in TCGA datasets, associating a subtype cluster B (PDGFRα, PDGFRβ, ACTA2, THY1, PDPN, FAP, and COL1A1) with worse prognosis and advanced stage, while subtype cluster A corresponded to better prognosis and an earlier stage ^36^. The conditioned CAFs showed a significantly higher CAF score than HUFs when scored for these seven genes (Figure 4). Mishra et al. ^37^ determined that by exposing human bone marrow-derived stem cells (hMSCs) to tumor conditioned media, the MSCs differentiate into a CAF-like phenotype expressing CAF-related genes. Similarly, when we used the top genes upregulated by the conditioned exposed hMSCs to assign a CAF score, KURA-CAFs and OC-CAFs expressed higher scores against HUFs (Figure 4), supporting the use of tumor conditioned media to reprogram stromal cells into CAFs. These outcomes together suggest that the generated conditioned CAFs resemble a CAF stromal-like subtype associated with worse outcomes in OC patients.

In terms of functional characterization of the CAFs, several avenues were addressed to provide a deep and thorough investigation. An accepted marker of fibroblast activation is collagen contractility ^66,67^. The contractile forces of activated fibroblasts are increased by signals from the ECM and α-SMA ^68^ and can be determined by a collagen gel contraction assay. Through this assay, our conditioned CAFs displayed a significant contraction of collagen type 1, which confirmed fibroblast activation (Figure 3). Another widely known trait of CAFs is their ability to induce tumor promotion and drug resistance in the TME ^4,69^. We confirmed these characteristics in our conditioned CAFs by performing co-culture experiments with cancer cells in 3D gel matrices and spheroids. In both experiments, our results show that when conditioned CAFs are co-cultured with OC, there is a significant increase in tumor growth and an increased potential to promote chemoresistance when compared to cancer alone and cancer co-cultured with normal fibroblasts (HUFs) (Figure 3). Because of CAFs tumor-promoting role in the TME, co-culturing CAFs with OC likely improved cancer cell attachment, growth, cell-cell interactions, and cell-ECM interactions ^70,71^. Furthermore, a higher ALDH expression in CAFs has been linked to increased stemness, where cancer-associated MSCs from human epithelial ovarian cancer compared to normal ovary-derived MSCs had a 2-fold increase in the percentage of ALDH+ cells ^72^, which was consistent with our findings (Figure 3). These results confirm that both characterized conditioned CAFs are activated fibroblasts recapitulating key functional CAF subtypes.

The ECM is significantly gaining attention as a key contributor to OC tumor progression and recurrence ^73,74^. To better study the ECM, the matrisome was established as an ensemble of over 1000 ECM-related genes that encode for two groups of ECM proteins, either matrisome-associated or core matrisome. ECM-associated proteins include ECM-affiliated proteins, ECM regulators, and secreted factors known or suspected to bind core ECM proteins, and the core matrisome includes glycoproteins, collagens, and proteoglycans. Stromal cells are the main contributor to ECM secretion and to matrisome expression. We identified that the reprogramming of HUFs into CAFs induces severe transcriptomic changes in the matrisome over both matrisome-associated and core matrisome categories. Figure 4 shows that many matrisome-related genes and collagens are upregulated by the CM-CAFs and downregulated by the HUFs, which supports the role conditioned CAFs play in ECM remodeling ^75,76^. Moreover, our data clearly implicated the conditioned CAFs with TGF-beta signaling, which is associated with tumor promotion, cancer cell invasion and migration, metastasis, and poor overall survival ^15,53,77^. Figure 4 shows upregulation of related TGF-beta signaling pathways SMAD and STAT in the conditioned CAFs ^78–80^. The immunomodulatory role of the conditioned CAFs was confirmed by the upregulation of pro-inflammatory genes and downregulation of anti-inflammatory genes (Figure 4). This coincides with previous research highlighting CAFs role in tumor-promoting inflammation ^11,81^. Of note, the anti-mullerian hormone (AMH) gene is also upregulated by the conditioned CAFs, which has been recently linked to the modulation of cancer-associated mesothelial cells ^47^. These transcriptomic findings confirm a CAF-like transcriptional signature expressed by the ovarian conditioned CAFs and their potential role in ECM remodeling within the ovarian microenvironment.

Retrieval experiments were performed to determine the stable reprogramming of normal fibroblasts into CAFs. CAFs removed from tumor conditioned media remained CAF-like phenotypically and functionally. We speculate that the OC cells reprogram the normal fibroblasts into CAFs through epigenetic changes driven by cytokines, exosomes (miRNAs), and/or cell-free DNA (cfDNA) present in the conditioned media ^82^. Albrengues et al. observed an epigenetic switch initiated by the cytokine LIF in three cancer types that sustained the pro-invasive phenotype of CAFs ^83^. In ovarian cancer, it was found that the downregulation of miR-214 and miR-31 partnered with the upregulation of miR-155 was responsible for the activation of normal fibroblasts into CAFs ^84^. Also, it has recently been demonstrated that three different exosomal miRNAs derived from triple-negative breast cancer activate stromal fibroblasts ^85^. And Filatova et al. found that cfDNA originating from tumors can penetrate recipient cells and increase oncogenicity ^86^. These previous studies suggest epigenetic changes and paracrine signaling as mediators of CAF reprogramming. Further studies will need to elucidate the mechanisms involved in the reprogramming of the generated conditioned CAFs.

Taken together, our study presents a reproducible, cost-effective, and clinically relevant translatable-based approach to reprogram normal uterine fibroblasts into ovarian cancer-associated fibroblasts using ovarian tumor-derived conditioned media. Two ovarian conditioned CAFs were phenotypically and functionally characterized demonstrating an activated fibroblast status and resembling a CAF subtype associated with worse prognosis in OC patients. Our results are expected to have an important positive impact because they will provide strong evidence for further development of therapeutics that possess potentiality and specificity towards CAF-mediated chemotherapy resistance in OC.

### Limitation of the study

Future investigations will need to validate the versatility of our approach and investigate other cell-type origins besides uterine fibroblasts. CAFs can be derived from many different cell types ^10^, thus new studies will need to investigate reprogramming with the tumor conditioned media of these other cell types, like MSCs, epithelial, and endothelial cells. Additionally, the believed origin for ovarian cancer is the fallopian tube ^87,88^, hence further studies should investigate fallopian tube fibroblasts instead of uterine fibroblasts and characterize the functional changes and differences of the conditioned CAFs deriving from different cells of origins. Uterine fibroblasts were selected due to their gynecologic origin and reliable availability through ATCC. It has been shown that CAFs originate from local fibroblasts in OC both in animal models and *in vitro* ^89,90^, and we further confirmed that uterine fibroblasts can be reprogrammed into ovarian CAFs.

## ACKNOWLEDGMENTS

This project used Sanford Research Histology and Imaging Core, the Functional Genomics and Bioinformatics Core, and the Flow Cytometry Core Facilities that are supported in part by a Center for Cancer Research CoBRE and Center for Pediatrics Research CoBRE grant from the National Institutes of Health (5P20GM103548 and 5P20GM103620). We want to thank Claire Evans and Kelly Graber from the Sanford Research Histology and Imaging Core, respectively; Malini Mukherjee from the Sanford Functional Genomics and Bioinformatics Core, and the Flow Cytometry Core Facilities for training and help in experimental set up. We also want to thank Mariah Hoffman for her help with the script for DESeq2 in R. We would also like to thank the team from the Animal Research facility at Sanford Research that helped with the *in vivo* experiment. Research reported in this publication was supported by the National Institute of General Medical Sciences of the National Institutes of Health under Award Numbers 5P20GM103548 and U54GM128729, and by National Cancer Institute of the National Institutes of Health under award number R21CA259158. The content is solely the responsibility of the authors and does not necessarily represent the official views of the National Institutes of Health. The PI (P.P) also acknowledge support from the Governor’s Research Center and South Dakota Board of Regent’s and the Haarberg 3D Center.

## AUTOR CONTRIBUTIONS

Conceptualization, P.P.; Methodology, H.A., S.P., K.C., and J.W.; Investigation, H.A., S.P.; Resources, H.A., S.P., M.J. and J.W.; Writing – Original Draft, H.A. and P.P.; Writing – Review & Editing, H.A. and P.P.; Supervision, P.P.; Funding acquisition, P.P.

## METHODS

### Reagents

Type I collagenase, thrombin, trans-4-(aminomethyl)cyclohexanecarboxylic acid (AMCHA), 10% neutral buffered formalin, and calcium chloride (CaCl_2_) were purchased from Sigma-Aldrich (Saint Louis, MO, USA). Lipophilic tracer 3,3′-dioctadecyloxacarbocyanine perchlorate (DiO, excitation, 488 nm; emission 525/50 nm) was purchased from Invitrogen (Carlsbad, CA, USA). Counting beads for flow cytometry experiments were purchased from Biolegend (424902, San Diego, CA, USA). The drug carboplatin (CARBO) used for drug resistance studies was purchased from MedKoo (Morrisville, NC, USA). For the contraction assay, rat tail collagen type I was purchased from R&D Systems (Minneapolis, MN, USA) and glacial acetic acid was purchased from Fisher Scientific (Hampton, NH, USA).

### Cell Lines

Primary human uterine fibroblasts (HUFs) were purchased from ATCC (Manassas, VA, USA). HUFs were grown in DMEM media containing 1ng/mL fibroblast growth factor (FGF), 10% fetal bovine serum (FBS, Gibco, Life Technologies, Grand Island, NY, USA), 5ug/mL insulin, 1% penicillin/streptomycin (Corning CellGro, Mediatech, Manassas, VA, USA). Human serous ovarian cancer cell line SKOV-3 and high-grade serous ovarian cancer cell line KURAMOCHI were gifts from Dr. Paola Vermeer (Sanford Research, Sioux Falls, SD, USA). OC cell lines were grown in ovarian media, a 1:1 ratio of DMEM and Ham’s F12 with the addition of 1% penicillin/streptomycin and 10% FBS. The primary OC-CAF cell line was obtained from Vitro Biopharma (Denver, CO, USA) and was grown in MSC-GRO Vitroplus media with 1% penicillin/streptomycin. Adipose-derived MSCs were purchased from Fisher Scientific and grown in MesenPRO media growth supplement with 2% MesenPRO growth supplement, 1% L-glutamine, and 1% penicillin/streptomycin. All cells were cultured at 37 °C, 5% CO_2_, and 21% O_2_; primary OC-CAF cells were also cultured in a hypoxic environment at 37 °C, 5% CO_2_ and 1.5% O_2_.

### Human Samples

Blood from healthy donors was collected at Sanford Biobank in Sioux Falls, SD. Informed consent was obtained from all subjects with approval from the Sanford Health Institutional Review Board and in accordance with the Declaration of Helsinki. Peripheral blood from healthy subjects was obtained through venipuncture and collected in whole blood collection tubes (BD Lavender K2-EDTA Vacutainer). Plasma was separated by centrifugation of the blood samples at 800 *x g* for 10 min, then a secondary spin at 400 × *g* for 10 min with plasma immediately aliquoted and stored at −80 °C.

### HUFs Reprogrammed into CAFs

For the reprogramming of HUFs into CAFs, tumor-derived conditioned media from both OC cell lines was collected, centrifuged, and filtered with syringes using a 0.2 um filter. Conditioned media was isolated from confluent OC cultures and kept at 4⁰C for no more than 1 week. Conditioned media was supplemented with 10% FBS at the time of culture. Two types of conditioned CAFs were generated, SKOV-3 conditioned CAFs (SKOV-CAFS), and KURAMOCHI conditioned CAFs (KURA-CAFs). Conditioned CAFs were assessed until passage 12, including retrieval, and at least 3 different lots of HUF cells were tested throughout experiments.

### Cell Expansion Analysis

Expansion of HUFs, conditioned CAFs, and primary OC-CAFs was assessed by plating 1.0 × 10^5 cells/mL in triplicates in 96-well flat bottom plates (Fisher Scientific) and growing for 7 days between cell passages before trypsinization, and finally counting cells by Countess II (Fisher Scientific). Proliferation of each cell type is reported as the expansion fold after 7 days for each passage.

### Cell Surface Marker Characterization

HUFs, conditioned CAFs, primary OC-CAFs, SKOV-3, KURAMOCHI, and adipose MSCs were phenotypically characterized by analyzing CAF markers (FAP, CD29, PDGFRα, α-SMA), stromal markers (CD90, CD73, vimentin), epithelial markers (EpCAM), and immune marker (CD45) (Table S1). 0.1 × 10^6^ cell/mL cells were stained with the appropriate antibodies and analyzed by flow cytometry (BD FACS Fortessa, BD Biosciences, San Jose, CA, USA). Fluorescence minus one (FMOs) samples were used as controls.

### Tumor Progression and Drug Resistance in Physiologically Relevant 3D Cultures

HUFs, conditioned CAFs, and primary OC-CAFs were embedded in a physiologically relevant three-dimensional (3D) culture system that recapitulates the complex biology of the TME and allows better cell-cell and cell-ECM interactions ^91–94^. We performed a co-culture of cancer and stromal cells to evaluate tumor promotion by the different stromal cell types. SKOV-3 and KURAMOCHI OC cells (0.3 × 10^5^ cell/mL) were prelabeled with DiO (10ug/mL) and incorporated into a 3D matrix either alone, or in a co-culture with HUFs, their respective conditioned CAFs, (SKOV-CAFs or KURA-CAFs), and primary OC-CAFs. The 3D cultures were established by mixing plasma, cells in ovarian media, a crosslinker, and a stabilizer in a 4:4:1:1 ratio, as previously described ^91–94^. After stabilization, ovarian media was added on top of the 3D cultures and renewed every 3 days. After 7 days of incubation, type I collagenase (20 mg/mL for 2 h at 37 °C) was used to enzymatically digest the 3D matrices. The isolated cells were stained for cell viability using a Live/Dead Blue cell stain (L34962, Thermo Fisher Scientific, Waltham, MA, USA) and blocked with 4% bovine serum albumin (BSA). Counting beads were added to each sample and a minimum 10,000 events were acquired by BD FACS Fortessa (BD Biosciences, San Jose, CA, USA) as previously described [21-24]. The data was analyzed with FlowJo program v10 (BD Biosciences, San Jose, CA, USA). Cancer cells were identified by gating high DiO signal. The effect of stromal cell types on drug resistance was also studied and carboplatin was given in order to recapitulate the standard of care treatment in OC. The same 3D matrix cultures were exposed to the drug carboplatin (30uM) against an untreated control for 7 days, and the same steps were followed as described above.

### Collagen Gel Contraction Assay

HUFs, conditioned CAFs, and primary OC-CAFs were incorporated into a 3D matrix made of rat tail type 1 collagen (R&D Systems) as previously described ^95^. A 3 mg/mL concentration of collagen was made by diluting rat tail type 1 collagen with 0.1% acetic acid. This collagen solution was then combined with cells in media suspension to form a collagen solution with a concentration of 1mg/mL. A predetermined volume of 1 M NaOH was quickly added and then the mixture was immediately plated in a 24-well flat bottom plate (Fisher Scientific) and allowed to solidify at room temperature before adding media and dissociating the gels from the sides of the well. The 3D matrices were incubated at 37°C for 24 hours while they were observed for contraction. Images were taken with the ChemiDoc MP Imaging System (Bio Rad, Hercules, CA, USA) at time intervals of 0 h and 24 h. ImageJ software (National Institutes of Health, Bethesda, MD, USA) was used to record the diameter and area change of the matrix images over a 24-hour period.

### Effect of Stromal Cell Types on Spheroid Formation

HUFs, conditioned CAFs, and primary OC-CAFs (0.42 × 10^5^ cell/mL) were co-cultured with either SKOV-3 or KURAMOCHI cells (0.14 × 10^5^ cell/mL), then plated on ultralow attachment 96-well microplates with clear bottoms (Corning, Corning, NY, USA). The spheroids were grown for 21 days and imaged every 7 days with Cytation 3 Imaging reader (Biotek, Winooski, VT, USA). ImageJ software was used to measure the lengths and areas of the spheroids to compare the formed spheroids over time. After 21 days, the spheroids were enzymatically digested with type 1 collagenase and stained with Live/Dead Blue cell stain, stromal marker CD90, and the ovarian cancer epithelial marker CD276 ((Table S1); then, counting beads were added, and the cells were analyzed with flow cytometry (BD FACS Fortessa). Results were analyzed in FlowJo program v10 to determine the composition of the different spheroid types.

### Cytokine Analysis

Cell-derived conditioned media and corresponding culture media from HUFs, conditioned CAFs, and primary OC-CAFs was functionally characterized by cytokine array using the C-Series Human Cytokine Antibody Array C5 (RayBiotech Inc., Norcross, GA, USA), following the instructions provided by the manufacturer. The LI-COR Odyssey (LI-COR Biosciences, Lincoln, NE, USA) device was used to capture images of the chemiluminescence signals of each of the membranes with a 5 min exposure time. The chemiluminescence signal intensity of each array dot was determined by subtracting the negative controls and then normalizing the positive controls in each membrane.

### Collagen Deposition by Immunohistochemistry (IHC)

3D matrices containing monocultures of HUFs and conditioned CAFs were fixed in 10% neutral buffered formalin and processed on a Leica 300 ASP tissue processor. 3D matrix slides were stained by IHC with α-collagen 1 diluted 1:50 (Table S1), with DAB as the chromogen and the slides were counterstained with hematoxylin. Omission of the primary antibody served as the negative control. The Aperio VERSA Bright Field Fluorescence & FISH Digital Pathology Scanner (Leica Biosystems, Buffalo Grove, IL, USA) was used to image the slides, which were then quantified for positivity (Npositive/Ntotal) to account for potential differences in collagen deposition in Aperio Imagescope (Leica Biosystems, Buffalo Grove, IL, USA).

### ALDEFLUOR Assay

ALDH activity was assessed in HUFs, conditioned CAFs, and primary OC-CAFs using the ALDEFLUOR assay kit (STEMCELL Technologies, Cambridge, MA, USA). The assay was performed per manufacturer’s protocol. 0.1 × 10^6^ cell/mL cells were resuspended in the Aldefluor assay buffer containing ALDH substrate (BODIPY-aminoacetaldehyde). The ALDH inhibitor diethylamino benzaldehyde (DEAB) was added to half the cells to provide a negative control. The cell suspensions were incubated at 37°C for 45 min, suspended in PBS 1x and counting beads, and analyzed to measure fluorescence by BD FACS Fortessa. The percentage of ALDEFLUOR-positive cells for each sample was gated separately and compared to the negative controls marked by the inhibitor DEAB in FlowJo program v10.

### Retrieval Experiments

Retrieval of tumor-derived conditioned media was also investigated, where conditioned CAFs were kept in culture with the corresponding tumor-derived conditioned media or switched to ovarian media, depriving them from the conditioned media exposure. The conditioned CAFs and retrieval CAFs were reassessed phenotypically and functionally through marker expression, collagen contractility, and tumor growth as described above for passages 1, 3, and 5 post-conditioned media retrieval.

### RNA-sequencing Analysis

RNA from the HUFs, conditioned CAFs, and primary OC-CAFs was sequenced. Cell pellets were flash frozen in liquid nitrogen and then RNA was extracted using QIAgen RNeasy mini columns according to manufacturer’s protocol ^96^. RNA was quantified using a NanoDrop spectrophotometer (Thermo Fisher Scientific, Waltham, MA, USA) and the Agilent 2100 Bioanalyzer (Santa Clara, CA, USA) was used to verify RNA quality under the support of the Functional Genomics and Bioinformatics Core at Sanford Research. Sequencing was performed by Novogene. FASTQ was used to check the read quality of the raw sequencing reads and STAR was used for mapping the reads to genome. In RStudio, the aligned read counts by gene were converted into a count matrix, which was then analyzed by DESeq2 (version 1.40.2) to identify differential expression by normalizing counts and producing differentially expressed genes (DEGs). DEGs were functionally analyzed using the clusterProfiler R package for gene ontology (GO) analysis. All volcano plots were built using the EnhancedVolcano R package and all heatmaps were built using the ComplexHeatmap R package. Using the RNA-sequencing results for HUFs, CM-CAFs, and OC-CAFs with the GSVA R package, an ECM score was produced using three gene sets from the GSEA website for the core, associated, and complete Naba matrisome and a CAF score was produced from gene sets used in previous studies: 24 carcinoma-associated fibroblast genes upregulated by MSCs grown in tumor conditioned media ^37^; 7 CAF markers used to identify two CAF subtypes ^36^; and 5 CAF markers used to identify four CAF subtypes ^38^. A dataset of OC patients from Tothill et al. ^35^ was obtained through accession GSE9899. The affy R package was used to read .CEL files containing affy IDs and export an expression data file where affy IDs were converted to Entrez gene symbols with the website DAVID. A mesenchymal score was produced with GSVA using all genes that were upregulated in the mesenchymal (C5) subtype that the paper identified, where we used only the top 10% of mesenchymal^high^ scoring samples (29 total). This data from the 29 mesenchymal patients was then merged and normalized in R with the RNA-sequencing results for normal HUFs, and then analyzed with DESeq2 to produce DEGs. DEG lists were annotated using the Matrisome AnalyzeR tool. The venn diagram R package was utilized to compare the overlapping DEGs among the datasets.

### Tumorigenic Potential of Conditioned CAFs in vivo

All *in vivo* experiments were conducted according to protocols approved by the Institutional Animal Care and Use Committee (IACUC) at Sanford Research. 0.3 × 10^6^ cell/mL of HUFs, conditioned CAFs, and primary OC-CAFs was injected into female, 6–10-week-old immunodeficient Swiss Webster mice intramuscularly in the flank, and tumor progression was evaluated by caliper for 6 weeks. Twice a week the mice were weighed, and tumor size was measured. The formula V (mm3) = 1/2 × length (mm) × width (*2) (mm) was used to determine tumor growth.

### Data Availability Statement

The datasets analyzed during the current study are available at the public Gene Expression Omnibus (GEO) database and are detailed in the methods section. Any other data can be available upon reasonable request to the corresponding author.

### Statistical Analysis

All *in vitro* experiments were repeated independently at least three times with triplicate samples per experiment unless otherwise stated. GraphPad Prism 5 (GraphPad Software Inc., La Jolla, CA, USA) was used for graphing results. We have presented the results as a mean ± a standard deviation value. Statistical significance was analyzed using Student’s t-test, one-way ANOVA, or two-way ANOVA, where a *p* value less than 0.05 was considered significant and an interquartile range (IQR) was used to remove outliers^91^. Unless stated otherwise, *p>0.05, **p>0.01, ***p>0.001, and ****p>0.0001.

