## Supplemental figures 1-3 and Table S1 for "Normal Uterine Fibroblast Are Reprogramed into Ovarian Cancer-Associated Fibroblasts by Ovarian Tumor-derived Conditioned Media"

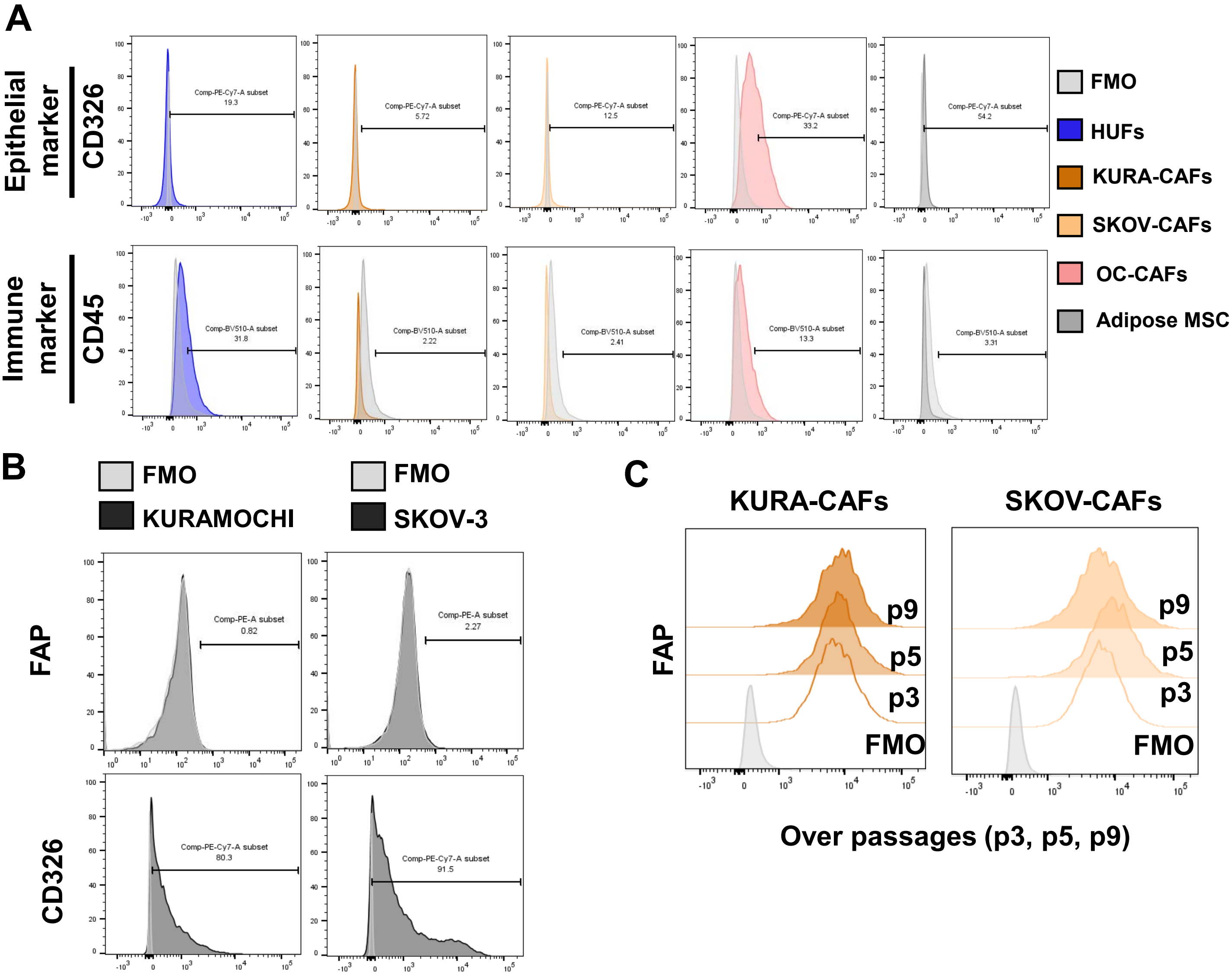


**Figure S1. Phenotypic marker expression, related to Figure 1** (A) Representative flow cytometry histograms of expression of epithelial markers (EpCAM/CD326), and immune markers (CD45) and respective fluorescence minus one (FMO) controls in HUFs, KURA-CAFs, SKOV-CAFs, OC-CAFs, and adipose-derived MSCs. (B) Representative flow cytometry histograms of expression of FAP and CD326 and respective FMO controls in KURAMOCHI and SKOV-3 OC cell lines. C) FAP expression over passages (p3-p9) for KURA-CAFs and SKOV-CAFs and corresponding FMO control

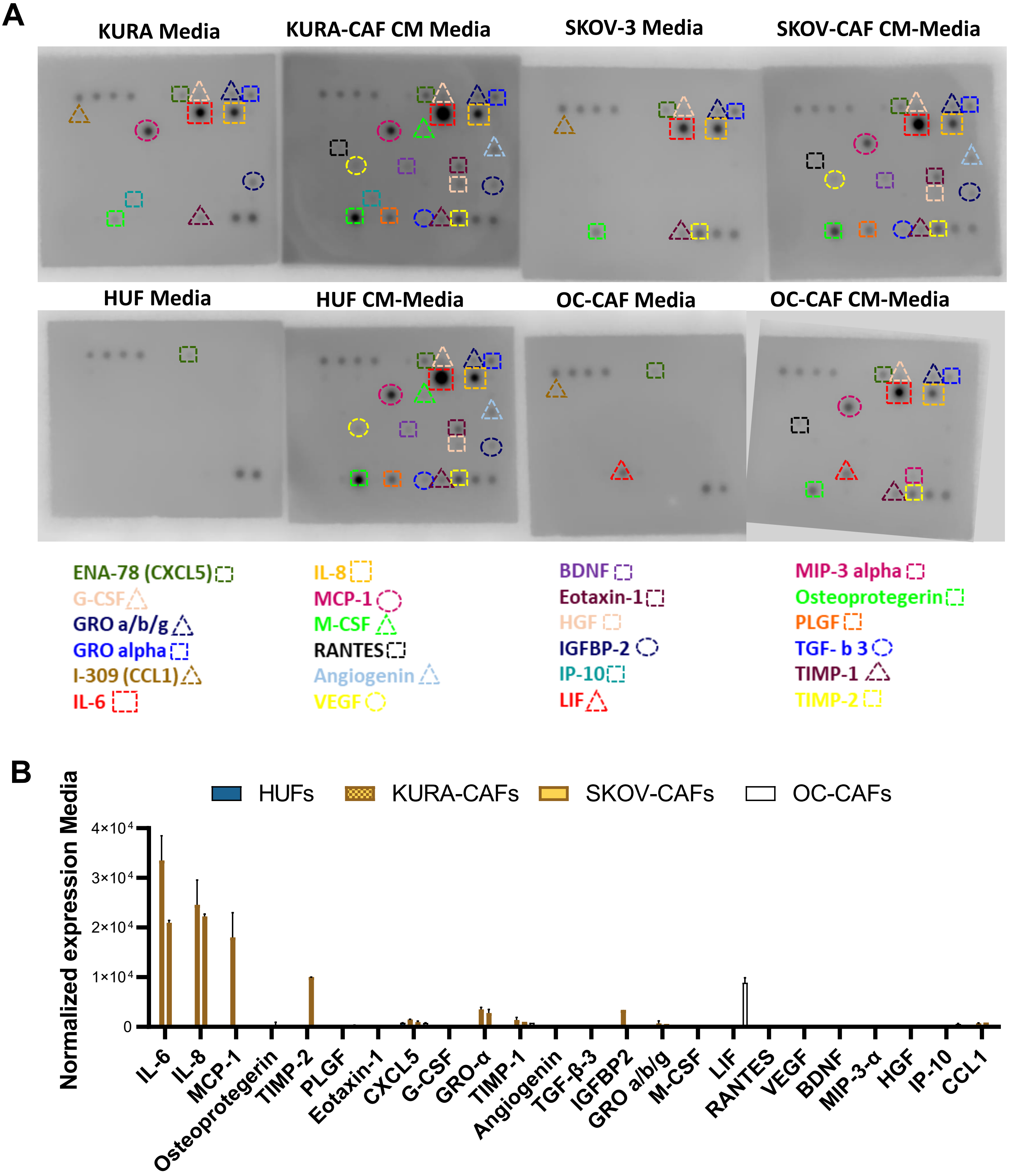


**Figure S2. Cytokine array, related to Figure 2** (A) Representative cytokine array image of cell-specific media only (Media) and media conditioned by cells cultured for 4 days (CM-Media) with expressed cytokine markers labeled by color. (B) Normalized expression of cytokines

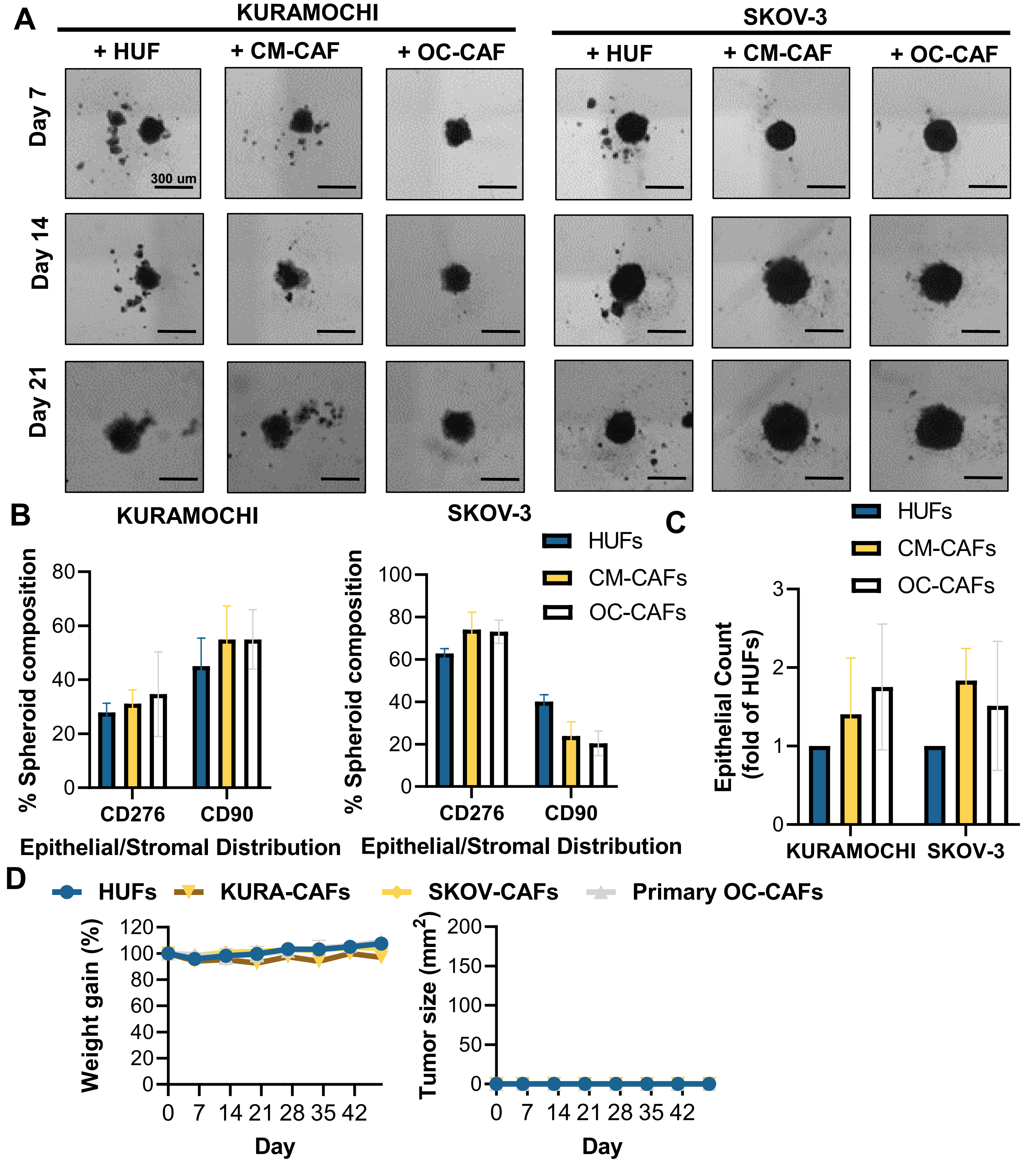


**Figure S3. Spheroid composition and *in vivo* tumorigenic potential, related to Figure 3.** (A) Representative Cytation3 images of KURAMOCHI and SKOV-3 cancer spheroids co-cultured with HUFs, CM-CAFs, and OC-CAFs, respectively on day 7, 14, and 21. Scale bar = 300μm. (B) Spheroid composition for stromal marker (CD90) and epithelial marker (CD276) representing the epithelial/stromal distribution in KURAMOCHI and SKOV-3 spheroids co-cultured with HUFs, CM-CAFs, and OC-CAFs, respectively, on day 21. (C) Epithelial count of KURAMOCHI and SKOV-3 spheroids co-cultured with HUFs, CM-CAFs, and OC-CAFs, respectively, on day 21. (D) Weight gain and tumor size of mice injected with HUFs, KURA-CAFs, SKOV-CAFs, and OC-CAFs, respectively, over 6 weeks (n = 3).

| **Table S1. List of antibodies used in the study** | | | |
|---|---|---|---|
| **Antibodies** | **Host** | **Vendor** | **Dilution** |
| PE-FAP | Mouse | R&D Systems-FAB3715P100 | 1:1 |
| PE-CD29 | Mouse | Biolegend-303004 | 1:10 |
| FITC-PDGFRα | Mouse | Fisher Scientific-PIMA528585 | 1:10 |
| APC-αSMA | Mouse | Fisher Scientific-501124530 | 1:200 |
| FITC-CD90 | Mouse | Biolegend-328108 | 1:10 |
| PerCP/Cy5.5-CD73 | Mouse | Biolegend-344014 | 1:10 |
| APC-Vimentin | Mouse | Biolegend-677807 | 1:10 |
| PE/Cy7-CD326/EpCAM | Mouse | Biolegend-324222 | 1:1 |
| BV510-CD45 | Mouse | Biolegend-304036 | 1:10 |
| BV605-CD276 | Mouse | BD Biosciences-743527 | 1:1 |
| Anti-COL1A1 | Rabbit | Sigma Aldrich-HPA011795 | 1:50 |
